# RevGraphVAMP: A protein molecular simulation analysis model combining graph convolutional neural networks and physical constraints

**DOI:** 10.1101/2024.03.11.584426

**Authors:** Ying Huang, Huiling Zhang, Zhenli Lin, Yanjie Wei, Wenhui Xi

## Abstract

Molecular simulation (MD) is an important research area in the field of life sciences, focusing on understanding the mechanisms of biomolecular interactions at atomic scales. Protein simulation, as a critical subfield of life science, has frequently adopted MD for implementation, where the trajectory data play an important role in drug discovery. With the advancement of high-performance computing and deep learning technology, machine-prediction of protein properties from enormous trajectory data becomes popular and critical, which puts challenges on how to extract useful data features from the complicated simulation data and reasonably reduce the dimensionality. At the same time, in order to better study the Protein system, it is necessary to provide a meaningful explanation of biological mechanism for dimensionality reduction. In order to address this issue, a new unsupervised model RevGraphVAMP is proposed to intelligently analyze the simulation trajectory. RevGraphVAMP is based on the Markov variation method (VAMP) and innovatively integrates graph convolutional neural networks and physical constraint optimization to improve the learning performance of the model. Besides, the attention mechanism is introduced to calculate the importance of protein molecules, leading to interpretation of molecular mechanism. Compared with other VAMPNets models, the new model presented in this paper has achieved the highest VAMP scores and better state transition prediction accuracy in two public datasets. Additionally, it has higher dimensionality reduction discrimination ability for different substates and provides interpretable results for protein structural characterization.

## 1. Introduction

Molecular Dynamics Simulation (MD) [1-3] is a critical methodology employed for understanding the structure and function of biological macromolecules. It encompasses a range of techniques, including ab initio calculations, semi-empirical quantum mechanical simulations, molecular dynamics simulations, multi-scale mixed simulations, coarse-grained simulations, and molecular docking[4]. By employing these approaches, researchers can gain profound insights into the intricate workings of biological systems at the molecular level. The results obtained from MD of proteins have proven to be invaluable in understanding various crucial biological processes. MD simulations have found applications in diverse areas such as disease mechanisms, drug development, and disease diagnosis within the realm of disease-related research[5, 6].

Biological macromolecules, like proteins and nucleic acids, are the core of biological activities, and proteins are important molecular substances in life systems, which play a vital role in the construction of life systems. The structure of a protein is determined by its primary sequence structure, and serves as the key to comprehending vital biological functions such as membrane transport, enzyme regulation mechanism, DNA transcription. By exploring the structure-function relationship of proteins at the atomic level, human understanding of the structure, morphology and folding process of proteins can be enhanced. For example, molecular simulation can be used to study the functional properties of target proteins and protein-protein interactions at the molecular level. On the one hand, it aids in uncovering the intricate pathogenic mechanisms of diseases, offering a detailed understanding at a finer-grained level, and on the other hand, it facilitates the screening and testing of drugs specifically targeting key proteins, enabling efficient identification and evaluation of potential therapeutic compounds.[7, 8]. All-atom molecular dynamics simulations offer a detailed investigation of functional conformational changes. However, due to the lack of protein structure data and conditional verification force fields, and the inherent sequence properties of numerical integration require femtosecond-level short time steps to maintain numerical stability, the timescales achievable in protein simulations (microseconds or less) still fall orders of magnitude short of capturing the timescales of functional conformational changes (milliseconds or longer). Markov state models(MSMs) [9] is a widely adopted approach for bridging this timescale gap, enabling the prediction and analysis of long-scale dynamical trajectories by leveraging a multitude of short MD simulations. At present, MSMs find extensive application in the study of global conformational changes, such as protein folding and dynamic changes of disordered proteins[10-13].

As the molecular simulation time and the number of protein molecules increase, the dynamic trajectory of proteins encompasses an expanding dimensionality and data scale. Consequently, the challenge lies in extracting meaningful data features from the intricate simulation data and reasonably reducing its dimensionality. Most traditional analysis methods rely on researchers’ expertise in molecular simulation and domain-specific knowledge, making it difficult to achieve automated and efficient analysis devoid of such reliance [14, 15]. In addition, for the data mining method itself, how to correlate and explain the biological mechanism is also an important issue, such as how to determine the core physical quantity, core conformation, and core region of the protein when determining the drug target. The development of deep learning technology has brought the dawn to solve the above problems [16-18]. The variational approach for Markov processes method(VAMP)[19] enables the optimization of latent space representations and linear dynamics models through unsupervised machine learning techniques. VAMPnets[20] is one of the first architectures to implement MSMs with deep learning based on the VAMP principle. It integrates the entire data processing pipeline into a single end-to-end framework, enabling dimensionality reduction and reducing reliance on prior knowledge. It has higher computational precision when studying protein folding problems. However, VAMPnets is strongly dependent on the data, and the performance of the simulation will be weakened when there are a large number of non-equilibrium samples in the molecular simulation trajectory. Therefore, in an effort to address the limitations of the original VAMPnets, Wei et al. [21] proposed the State-free Reversible VAMPnets (SRV) model. This approach achieves detailed balancing by transforming the lag covariance matrix into a symmetric matrix, and applying its results to an enhanced sampling algorithm for CVs, facilitates new procedures for the analysis of protein trajectories. Wu et al. [22] introduced additional constraint variables to impose reversibility, developed a VAMPnets model with physical constraints, and achieved stateless reversible optimization. This method was applied to the analysis of disordered proteins by Thomacs et al. [23], which once again demonstrated the effectiveness of deep learning methods for analyzing protein properties with little prior knowledge. However, the aforementioned network models rely on all simple fully connected networks, which exhibit limitations in effectively modeling complex protein structures and fail to consider the analytical nature of molecular mechanisms. In recent years, graph neural networks have played an important role in the fields of protein structure prediction, protein molecular dynamics, and protein design[24-26]. GDyNets[27] improved the VAMPnets model based on graph neural network optimization and applied it to the research of materials field and showed high performance, but this improved method has not been applied to protein analysis. GraphVAMPNet[28] uses the graph convolutional neural network SchNet[29] to model protein molecules, builds a new VAMPNet framework to improve network performance in multiple protein folding data. Notably, GraphVAMPNet offers interpretability by incorporating an attention mechanism, allowing for the visualization of intermolecular interaction strengths and the contribution of individual molecules to the microstate. However, GraphVAMPNet only modifies the original VANMPNet network without adding physical constraints and reversible optimization, which makes the model in applied to the analysis of data simulated by multiple non-equilibrium distributions can be problematic.

The objective of this research paper is to propose a novel deep learning model that facilitates efficient data mining and analysis of molecular simulation results. The primary goals are to reduce data dimensionality in a biologically or physically constrained manner, address interpretability challenges, and offer new avenues for drug-aided design. To achieve this, the paper introduces the RevGraphVAMP model, which effectively captures relevant structural features and collective variables to describe local functional conformational changes. This model combines graph convolutional neural networks and physical constraint terms, resulting in improved learning performance. The inclusion of physical constraints enables the model to handle non-equilibrium molecular simulation trajectory data, while the attention mechanism effectively models the significance of protein molecules, enhancing interpretability of the biological mechanisms underlying the model. The RevGraphVAMP model outperforms similar VAMPnets models in terms of prediction scores and overall performance on two datasets, providing compelling data-driven evidence for protein drug target analysis through interpretable results.

## 2. Materials & methods

### 2.1. Datasets and data preparation

#### 2.1.1 Datasets

The protein trajectory data generated by the molecular dynamics simulation, each frame of the trajectory is the three-dimensional coordinate information of the protein molecule at different times. In this study, the analyzed data primarily consists of trajectory information. Two public data sets which are also used in previous VAMPnet model were utilized: the alanine dipeptide dataset[30] and Aβ_42_ polypeptide dataset[23]. The alanine dipeptide dataset contains three trajectories with a simulation time of 250ns. Each trajectory includes the three-dimensional coordinate positions of 10 atoms, 250,000 frames of structural information, and the sampling time interval of each frame is 1ps. The structure is shown in Figure S1. The Aβ_42_ polypeptide dataset includes 5,119 protein trajectories, encompassing a total of 1,259,172 frames. In this simulation, the sampling interval of each frame is 250ps, and the total trajectory length is about 315*μ*s. The trajectory is generated by Gromacs MD and FAST. The protein includes 42 amino acids, and the structure is shown in Figure S2 shown. The reason for selecting Aβ_42_ data is based on the following considerations: (1) Aβ_42_ protein is related to Alzheimer’s disease[31], a significant and incurable condition in developed countries. Analyzing the properties of this protein holds crucial societal implications. (2) The protein dataset contains longer simulation trajectories, more protein molecules, and multiple short trajectories. Analysis of this protein can verify our The performance and application value of the model in larger-scale molecular simulations. (3) The protein exhibits complex structural changes and functions, with a certain degree of disorder[32]. Analyzing this protein presents a major challenge given its unique characteristics.

#### 2.1.2 Data preparation and preprocessing

To better capture the characteristics of proteins, we employ a graph data structure for protein representation. In the graph data structure, denoted as G = (V,E), V represents the characteristics of N nodes *V =* {*𝒱*_*1*,_ …, *𝒱*_*N*_}, while E represents the set of edges that capture the relationships between nodes E= {*e*_*1*,_ …*e*_*M*_}. For each protein structure, atoms or amino acids are selected as nodes, and the node embeddings obtained through random initialization or one-hot encoding. To establish the edges, the nearest m atoms or amino acids are selected for each node, and the edge embedding values are determined based on the Gaussian-expanded distance obtained from the following formula:

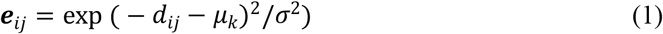

where *d*_*ij*_ is the distance between atoms *i* and *j*, and *σ* denotes the step size of the gaussian filter, *μ*_*k*_ = *d*_*min*_ + *k* × *σ* Å, *k* = 0,1, …, *n, n =* (*d*_*max*_ − *d*_*min*_)/ *σ, d*_*max*_ and *d*_*min*_ are the maximum and minimum distances for constructing Gaussian kernel expanded edge features, respectively.

After the above calculation, the distance *d*_*ij*_ between atoms *i* and *j* finally becomes an n-dimensional Gaussian expansion vector ***e***_*i*j_.

During the process of feature extraction, the polyalanine polypeptide dataset specifically chooses 10 atoms located on the main chain of amino acid residues as nodes. For each atom, its nearest neighbor atom is selected to calculate the edge embedding, thereby establishing the graph input. Similarly, the Aβ_42_ polypeptide dataset selects 42 amino acids as nodes, and the nearest neighbor amino acid is chosen for each amino acid to compute the edge embedding, resulting in the construction of the graph input.

### 2.2. The architecture of RevGraphVAMP

#### 2.2.1 The overall framework of RevGraphVAMP

The newly proposed RevGraphVAMP overall network model in this paper is shown in Figure 1. The idea of network analysis is based on the VAMP method. The overall architecture continues the end-to-end dimensionality reduction method of VAMPNets, integrates the graph convolutional network and physical constraints to increase the performance of the model, and increases the interpretability of the model through the attention mechanism network. The network input includes two protein conformations of time *t* and lag time *t* + *τ*. The corresponding graph representation is established after feature extraction of the protein data, which is input to the graph convolutional neural network model in the graph, and finally the dimensionality reduction transformation representation *χ*(*x*_*t*_) and *χ*(*x*_*t*+*τ*_) of the protein is obtained. During the training process, physical constraints are added through the U and S modules, and the entire network parameters are optimized by maximizing the VAMP score, and the optimal transformation *χ*(*x*) and Markov transition matrix are obtained.

**Figure 1.**
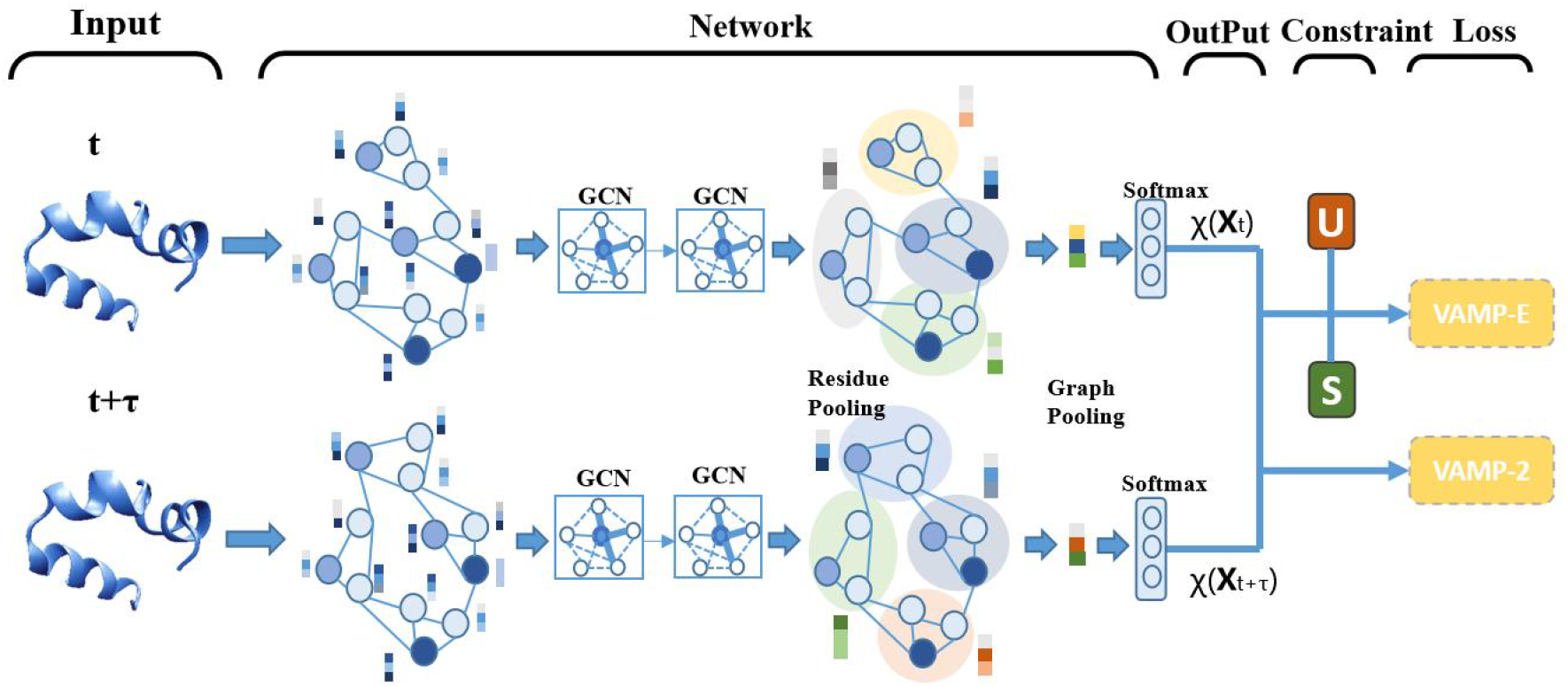
Network model architecture of RevGraphVAM

#### 2.2.2 Variational approach for Markov processes

Markov models estimate kinetic processes through transition densities. The transition density is the probability density of a transition to state y at time *t* + *τ* after a lag time *τ* or a given system in state x at time *t*:

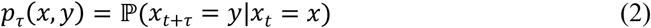

VAMP allow to find optimal feature maps and optimal dynamical Markov models from a given time series data. The core of the VAMP method is that the top singular component of the Koopman operator can obtain the best linear model. In Koopman theory, the Markov dynamics at lag time *τ* are approximated by a linear model of the form:

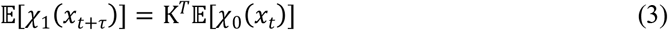

where *χ*_0_ (*x*) = (*χ*_01_ (*x*), …*χ*_0*m*_ (*x*))^*T*^ and *χ*_1_ (*x*) = (*χ*_11_ (*x*), …*χ*_1*m*_ (*x*))^*T*^ is the feature transformation, Convert the conformation x to a latent space representation that is linear in dynamics m-dimensionality, usually the space of slow transitions or rare events. K is the transition matrix, and 𝔼 represents the expected value changing with time, which explains the randomness in dynamics. The approximation error of the Koopman operator can be minimized by setting *χ*_0_ and *χ*_1_ as the upper-left and upper-right singular functions of the Koopman operator. The above equation minimizes the optimal matrix K of the regression error as:

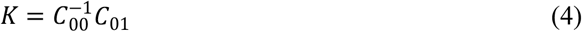

where the mean-free covariance matrix for the data transformation is defined as:

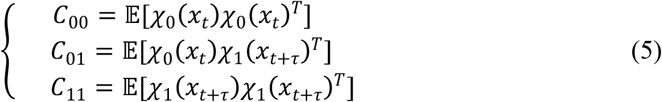

#### 2.2.3 Graph convolutional network of GASchnet

After careful analysis and testing, we have selected the optimized graph convolutional neural network component based on SchNet [29]from the GraphVAMPnet [28] proposed by Ghorbani et al. We refer to this component as GASchNet for brevity. GASchNet is primarily responsible for obtaining the protein trajectory’s feature transformation *χ*(*x*), and the network architecture details are shown in Figure 2.

**Figure 2.**
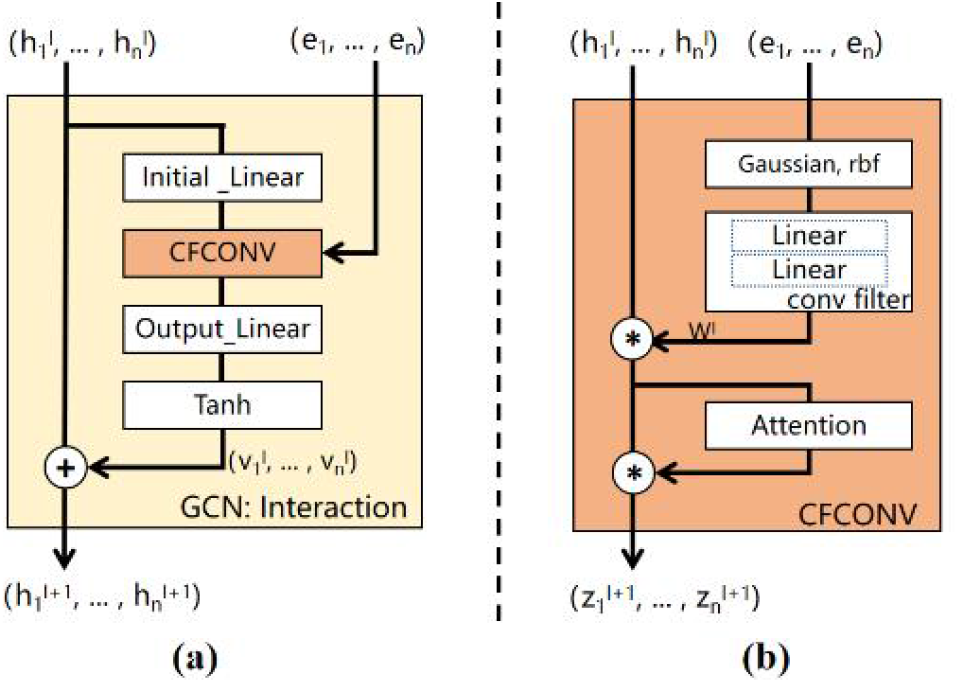
GASchNet graph convolution network structure

In GASchNet, protein features are extracted from graph data, and the model learns feature vectors on nodes in the graph. The node features 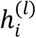 of the first round are given by node embedding, which are composed of trainable shared vectors with *d*_*h*_ dimensions. The initial value of node embedding 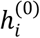 is obtained from the trajectory data according to the data processing method in Section 2.1.2. Node embeddings are updated in multiple graph convolutional interaction layers (GCN-Interaction), each of which contains continuous convolutions (CFCONV) between nodes. The edge embedding is calculated by the Gaussian distribution in the previous section, and then these edges are input into a conv filter w, which maps ***e***_*ij*_ to a *d*_*h*_ dimensional filter. This filter is then computed as a graph convolution to finally generate a new node embedding:

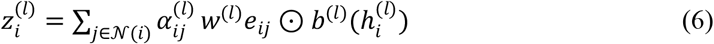

where *w* and *b* are both trainable functions, ⊙ is element-wise multiplication, 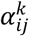 is the attention parameter, which will be explained later. The summed part of the formula means that the embedding update of node *i* in the network needs to include the embeddings of all neighboring nodes of *i* in graph 𝒩. This calculation method calculates all atomic interactions in the network through multiple interaction layers, so the network model can express complex many-body interactions. At the same time, this computing method uses node embeddings to represent the three-dimensional coordinates of molecules, and the convolution filter depends on the distance between molecules, thus maintaining the desired invariance in the system.

In addition, to increase the interpretability of the model, an attention layer[28] is added to the network, which learns the importance of edge embeddings for updating node embeddings, reflecting the interaction between a node and its neighbors. The analysis of attention weights can improve the interpretability of the model and establish the relationship with biological mechanisms. Note that the weights *α*_*ij*_ are learned between node embeddings and neighboring nodes using a Softmax function:

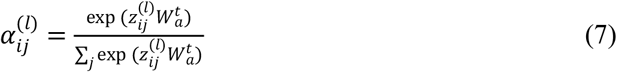

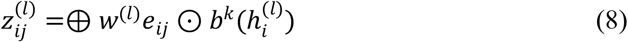

where 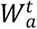 indicates the trainable parameters of the attention layer, 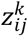 is the intermediate value of node *i* and *j* embedded in the filter generation network *w* and *b* calculated in the previous section, ⊙ indicates element-level multiplication, and ⊕ indicates the characteristics of adjacent nodes connect.

In each layer, node embeddings are updated in interactive layers, which can include a residual block to mitigate the issue of vanishing gradients, as commonly done in deep neural networks:

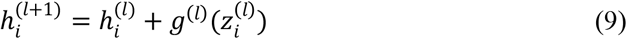

where *g*^(*l*)^ is a trainable function, including a linear layer and a nonlinear layer, where the nonlinear layer uses relu function.

The output of the final GASchNet interaction layer is fed to an atomically fully connected network. The node embeddings finally learned after multiple GASchNet interaction layers are input to the pooling layer, and the embeddings of all nodes are averaged to generate a graph embedding matrix for the protein structure of each time step *t*:

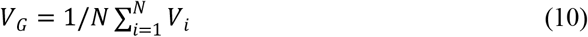

The model embeds the graph into a Softmax layer to obtain the final output, for all output *i* with ∑_*i*_ χ_*i*_ (*x*) = 1 for all χ_*i*_ (*x*) ≥ 0. so the value χ_*i*_ (*x*) of output node *i* can be expressed as the probability that the protein structure belongs to substate *i* at the current time *t*. If the network results are used for cluster analysis, then the substate to which the protein structure t belongs is the category corresponding to the maximum value of the output node. If the model is used for dimensionality reduction, the final output of the model is the dimensionality reduction result of the current protein structure through the χ(*x*) function.

#### 2.2.4 Physical constraints

To increase the model’s ability to analyze short trajectories from multiple non-equilibrium distributions, physical constraints need to be incorporated into the transition matrix K, in particular reversibility and randomness. After our analysis and testing, we choose the algorithm proposed by Wu et al. [22] to model reversibility and equilibrium state by introducing two new constraints S and u. After adding physical constraints, the transition probability density equation can be changed to:

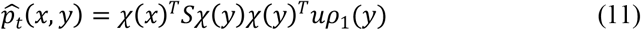

where S and *u* are trainable variables, *u* weights the empirical distribution as the equilibrium distribution of the system, and assumes that the system has a unique equilibrium distribution *μ* (*x*) = *μ*^*T*^χ (*x*) *ρ*(*x*)*.μ*(*x*) is the equilibrium distribution of the system state, *μ* (*x*)*p*_*τ*_(*x,y*)= *μ*(*y*)*p*_*τ*_ (*y,x*)

*ρ*_*1*_ represents the empirical distribution of *x*_*t*+*τ*_ in all transformations(*x*_*t*_, *x*_*t*+*τ*_). Therefore, the Koopman matrix K can be approximated as:

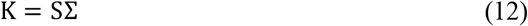

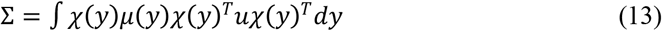

At this time, the Markov process defined by 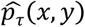 is a reversible Markov process with a stationary distribution *μ*(*x*).

The constraint part of the model is two constraint modules S and U, which generate parameters S and u to calculate the loss function. For the calculation of parameter u, the calculation is completed through a trainable weight *W*^*u*^:

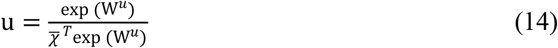

For the calculation of the parameter S, it is also realized through a trainable weigh *W*^*s*^:

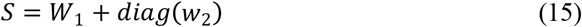

Among them, *W*_*1*_ is obtained from the trainable weight to ensure symmetry and non-negativity, *𝒲*_*2*_ changes the diagonal elements to ensure that the normalization *SC*_*11*_*u = S*_*𝒱*_= *1* does not affect the symmetry:

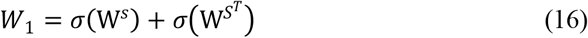

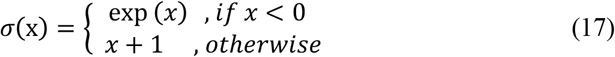

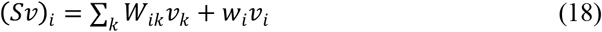

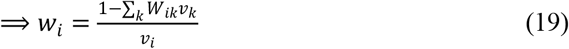

#### 2.2.5 Loss function

The loss function part of the model is divided into VAMP-E and VAMP-2 scores [19]. As mentioned above, the approximation error of the Koopman operator can be reduced by setting χ_*0*_ and χ_*1*_ as the upper left and upper right singular functions of the Koopman operator. Minimization, VAMP-E and VAMP-2 in the loss function are two variational scores, using the variational description of the singular component to measure the similarity between the estimated singular function and the true function. In model training, by maximizing the score Realize the training of model parameters, where VAMP-E is the main loss function, and the VAMP-E score is expressed as:

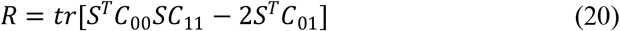

The VAMP-2 score is expressed as:

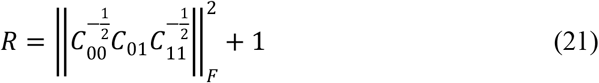

### 2.3. Training and inference

The training of the model is carried out as follows:

#### Algorithm 1 Model training strategy

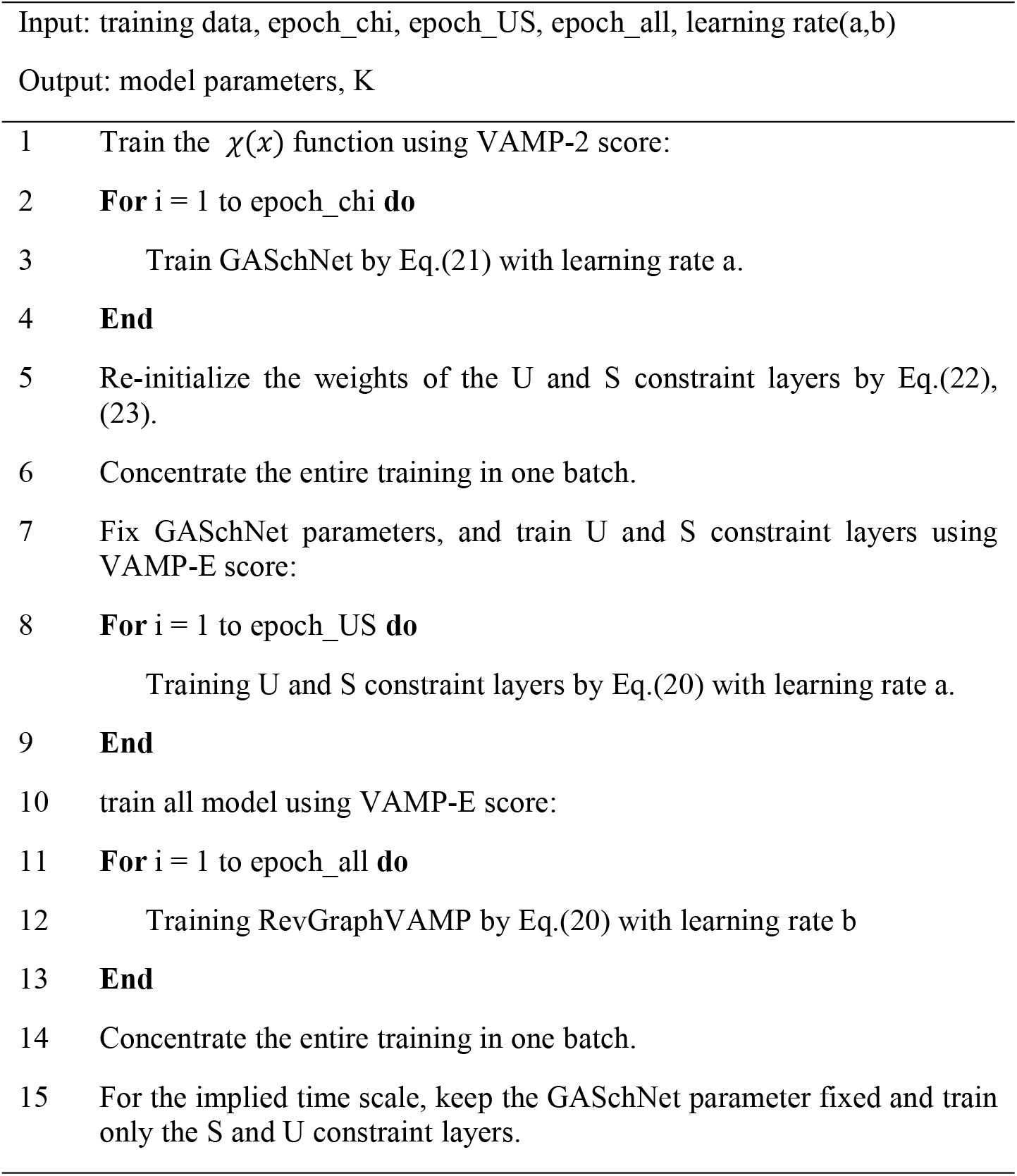

In step 5 of the above training process, the optimal initial value of the trainable parameter *W*^*u*^ *is updated as:*

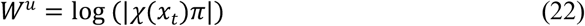

where π is the normalized stationary distribution of the characteristic values of the transition matrix K.

The optimal initialization value of the trainable parameter *W*^*s*^ is:

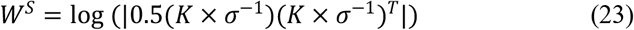

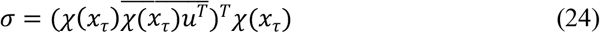

### 2.4. Model setting and evaluation metrics

The RevGraphVAMP model is based on the Markov model, so it needs to satisfy the Chapman−Kolmogorov (CK) equation:

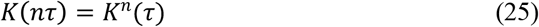

for all *n* ≥ 1, *K*(*nτ*) and *K*(*τ*) represent the models estimated in the lag time of n*τ and τrespectively*.

In order to select a suitable dynamical model, the test method is further developed based on Eq. (25), and the eigenvalue decomposition of each estimated Koopman matrix of the Markov state model is performed according to *K*(*τ*) = *r*_*i*_*λ*_*i*_ (*τ*), and compute the implied time scale [33]as a function of the lag time:

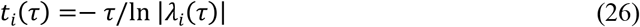

where *λ*_*i*_ (*τ*) is the eigenvalue of the Koopman matrix established on a lag time *τ*. Under this *τ* value, it is guaranteed that the implied time scale *t*_*i*_ (*τ*) is approximately constant in *τ*. Therefore, when verifying the correctness, it is necessary to select the minimum lag time *τ* when the implicit time scale *t*_*i*_ (*τ*) is approximately constant in *τ*, and after selecting the lag time *τ*, check whether the CK test (Eq.28) is statistically invariant within the range of certainty.

The implementation of all networks is based on Pytorch and Deeptime[34] software with NVIDIA GeForce GTX 3090. Pyemma[35] is used to draw the free energy landscape. The hyperparameters used in model training are shown in Table 1, using the mentioned in Section 2.3 For the training strategy, all models use the Adam optimizer, and the loss function uses VAMP-2 and VAMP-E. The data set is divided into a training set and a verification set according to 7:3.

**Table 1.**
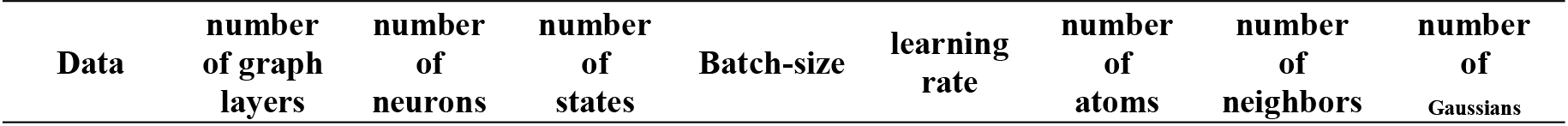
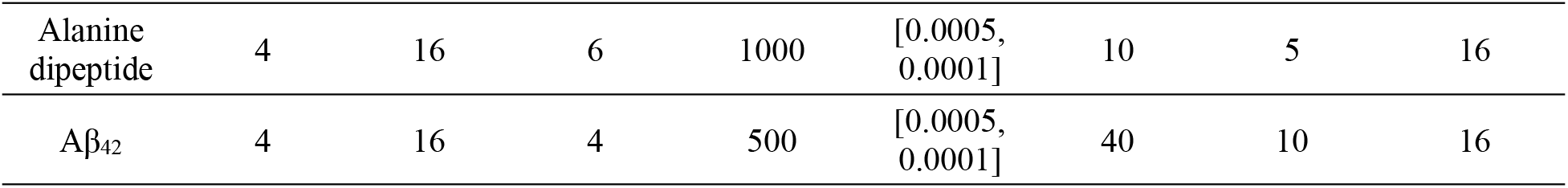
Hyperparameters of model training.

## 3. Results and discussion

In order to verify the effect of model optimization, we compared two classic VAMPnets networks [20, 23]and GraphVAMPNet’s GCN network[28] in previous studies, as shown in Table S1. At the same time, we also implemented a network based on The graph network (MAGNN) of the multi-head attention mechanism [36]is used to compare the effect of the model. This network uses the same feature input as RevGraphVAMP, uses the edge embedding as the key vector K and the value vector V, and the node embedding as the query vector Q. Update the node embedding to get the final graph embedding.

### 3.1. Analysis results of Alanine dipeptide data set

#### 3.1.1. Performance verification

In the Alanine dipeptide data set, the network is finally divided into 6 states. The distribution of each state is presented in Table S2, which is basically consistent with the model state division results in the original paper of VAMPnets. When comparing the VAMP scores with other models (Table 2), the model employing the graph convolutional network demonstrates higher VAMP-2 and VAMP-E scores compared to VAMPnets implemented using fully connected neural networks.

**Table 2.** Comparison of alanine dipeptide VAMP scores.

| Model | Alanine dipeptide | | Aβ<sub>42</sub> | |
| --- | --- | --- | --- | --- |
|  | VAMP-2 | VAMP-E | VAMP-2 | VAMP-E |
| VAMPnets | 4.32±0.23 | 4.35±0.02 | 3.98±0.008 | 3.93±0.050 |
| GCN-VAMP | 4.39±0.05 | 4.36±0.05 | 3.99±0.008 | 3.97±0.060 |
| MAGNN-VAMP | 4.41±0.03 | 4.36±0.03 | 3.97±0.001 | 3.95±0.002 |
| RevGraphVAMP | **4.41±0.01** | **4.38±0.01** | **3.99±0.002** | **3.99±0.003** |

This suggests that the graph convolution-based network is more effective in capturing data characteristics. Among them, the RevGraphVAMP model presented in this paper achieves the highest score, indicating that the model’s enhancements contribute to improved network accuracy.

The model presented in this paper demonstrates excellent performance in the CK test. The implicit time function of the trained model is depicted in Figure 3 a, with a selected lag time of τ=20 ps. The CK test results, illustrated in Figure 3b, exhibit a close alignment between the model’s predictions and the observed values for each of the six states. Notably, the transition curves for the first five states exhibit complete consistency, indicating a high accuracy of the Koopman matrix derived by the model. These results are consistent across multiple tests, as depicted in Figure 3. Furthermore, the small light blue confidence interval in Figure 3b signifies the model’s strong convergence.

**Figure 3.**
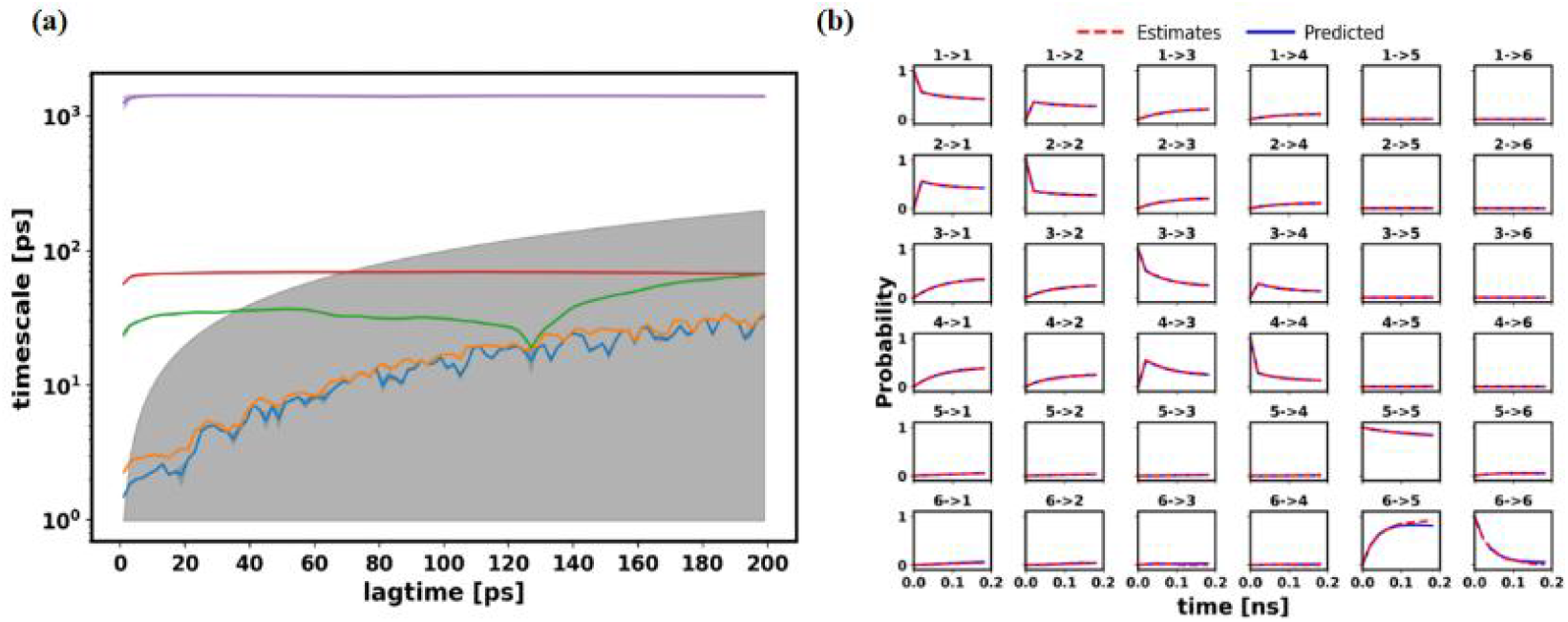
Correctness verification of alanine dipeptide. (a) Implied timescale (ITS), The light color part represents the confidence interval and the gray part represents the original time interval. (b) CK test results of 6 states,each subgraph represents the probability of transition from one substate to another at different times. The blue solid line is the predicted value K^n^(τ) of the transfer probability under this time, the red part is the estimated value K(nτ) calculated by using the K matrix according to Eq.(28), the light blue part is the confidence interval of multiple tests

According to the existing protein secondary structure mechanism theory, the classification of different dipeptide structures is typically based on the dihedral angles (referred to as Φ and Ψ) of the two peptide planes on the amino acid backbone. In our analysis, the two dihedral angles of the conformation of each protein in the trajectory are projected onto a two-dimensional plane, as illustrated in Figure 4. The heat map demonstrates that the model presented in this paper (Figure 4b) effectively distinguishes the characteristic structures, and the visualization of the model’s classification results on the Laplace diagram aligns with the conventional understanding of protein conformation distribution. This again proves that the model can effectively mine structural features Ability.

**Figure 4.**
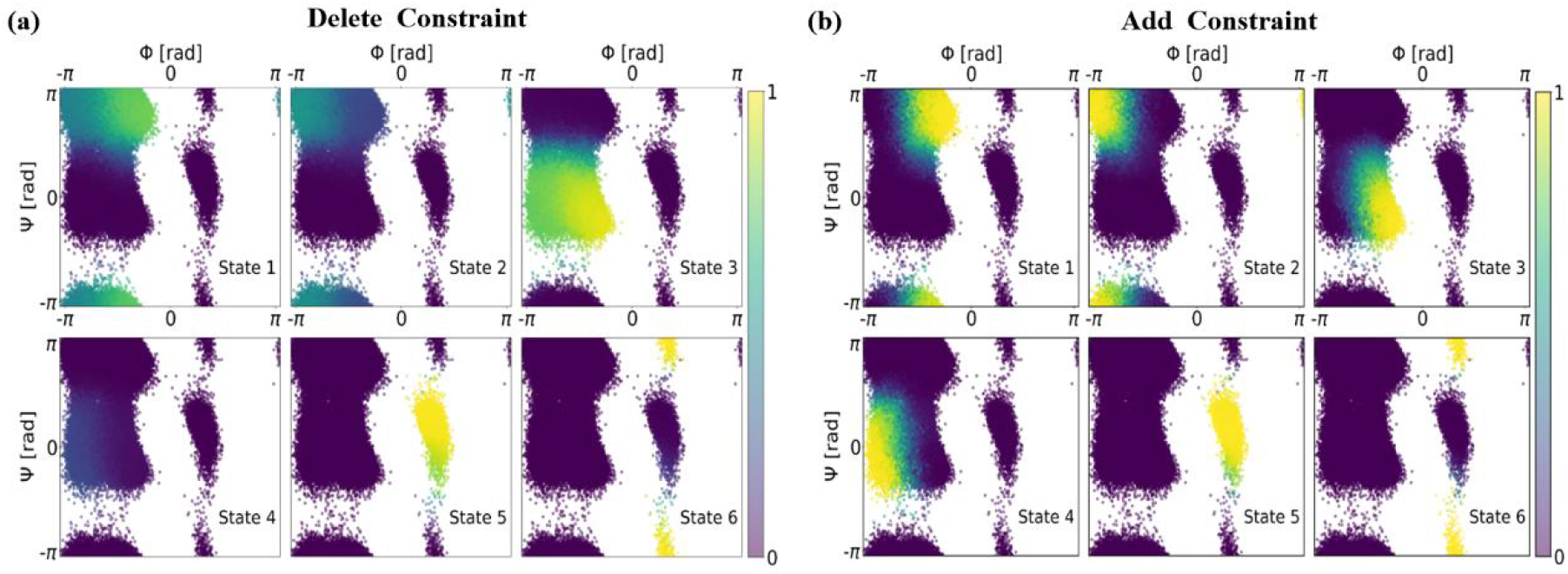
Angular heat map of six states divided by the model, plotted as a function of Φ and Ψ. (a) The result of 6 states with no physical constraints, (b) the result of 6 states add physical constraints. The colors correspond to the values of the respective output layers, indicating the probability of belonging to the associated metastable state

The physical constraints added in this paper enable the model to effectively differentiate between different metastable structures of dipeptides. In the two-dimensional projection heat map of dipeptide dihedral angle, when compared with the original unconstrained results (Fig. 4a), the state output value of this model in state 1-4 exhibit a clearer distribution in the figure. It demonstrates enhanced performance, particularly in distinguishing typical structures (indicated by brightly colored intervals), thereby indicating improved convergence and predictive capabilities of the model. The representative structure of the protein selected according to the highest score is shown in Figure 5a. The Φ and Ψ angles presented in the representative structure correspond to the values of the yellowest part of the heat map, which further verifies the correctness of this model.

**Figure 5.**
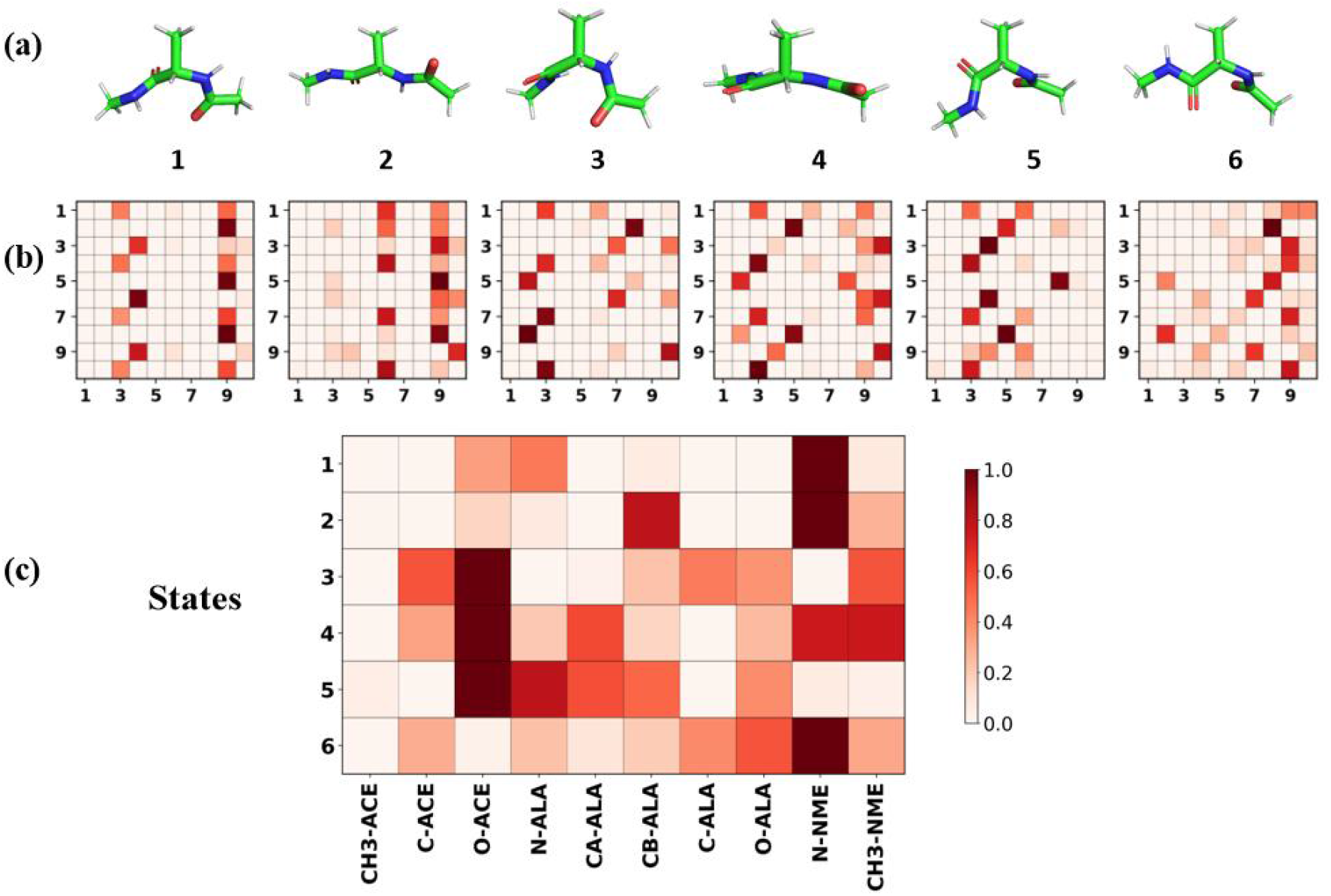
Dipeptide visualization results. (a) Typical conformation of six states, (b) average attention weights of 10 intermolecular interaction for dipeptides in each state, and (c) average attention weights of 10 atoms in different states. Darker red indicate that interactions between pairs of atoms are more important for the structural stability of the state. And data are derived from structures that exceed the 95% confidence threshold.

#### 3.1.2. Interpretability

In order to verify the interpretability of this model, we visualized the attention parameters of the final training of the model, and the results are presented in Figure 5 b and c. We selected 10 atoms on the main chain of amino acid residues and visualized the interaction strengths among these atoms for each state from 1 to 6. Notably, in state 1, nitrogen atom N-NME No. 9 demonstrates strong interactions with multiple atoms. The dihedral angle distribution and representative structure of state 2 (as shown in Figure 4) confirm that state 2 primarily characterizes the beta sheet structure, aligning with conventional understanding. Furthermore, in state 3, the oxygen atom O-ACE No. 3 emerges as particularly important, which is consistent with the alpha-helix structure depicted in Figure 4. Figure 5c provides additional insight by presenting the summed and averaged interaction strengths of the 10 atoms across all states. As observed, the oxygen atom O-ACE No. 3 and nitrogen atom N-NME No. 9 exhibit darker colors, indicating their significant contributions to the dipeptide structure. The nitrogen-oxygen atom pair is the key atom that forms the interaction between the peptide planes, which is consistent with the results of the attention mechanism in this paper. This outcome further emphasizes the utility of the attention mechanism in extracting and elucidating the molecular mechanisms of proteins. It offers researchers in related fields an end-to-end molecular mechanism analysis tool and data-driven mining methods.

### 3.2. Analysis result of Aβ_42_ data set

#### 3.2.1. Performance verification

Based on the training VAMP score, the graph neural network model proposed in this paper demonstrates higher accuracy. To validate its performance, a subset of 1024 randomly selected trajectories, totaling 273,715 protein structures, was used for verification. As shown in Table 2, the RevGraphVAMP model yields the best results. Its VAMP-E score surpasses the standard VAMPnets by 6%, the MAGNN-VAMP model by 4%, and the GCN-VAMP model by 2%, showcasing the model’s improvement in accuracy. Furthermore, through multiple tests, the SchNet-based graph convolutional network model presented in this paper exhibits better convergence, with a test score error of less than 1%. We implemented the standard VAMPnets with a network size consistent with Thomas’ article [23] in 2021, containing 464,646 parameters. In contrast, the model implemented in this paper consists of only 6,357 parameters. This significant reduction in the network model’s size effectively illustrates the advantages of utilizing graph convolutional neural networks.

In order to further compare the improvement of the performance of the graph neural network in this paper, according to the number of divided states as 4/5/6, we conduct the CK test on the classic VAMPnets and the K matrix trained by RevGraphVAMP in this paper. The test results, depicted in Figure S3, reveal that the predictions of the model in this paper closely align with the estimated data for the four divided states. Moreover, the Koopman matrix obtained by the model exhibits a high time delay and a smaller error interval, indicating superior convergence and enhanced predictive performance. At the same time, the transition error and confidence interval of the model in dividing 4-6 sub-states are observed to be smaller than those of the standard VAMPnets. This finding highlights the superior timeliness and convergence of the transition matrix trained using the graph neural network model proposed in this paper, thus validating the efficacy of the model framework.

After the previous test, the data is divided into 4 seed states with the best transfer performance. he model was then trained using the entire dataset to classify the data into these 4 states, resulting in the following probabilities: 52.3%, 26.7%, 10.7%, 10.2%. To further assess the accuracy of the predictions, the transition matrix K is decomposed into eigenvalues, and the implied time scale is calculated as a function of lag time. As shown in Figure 6, the time scales implied by the three eigenvalues all exceed original lag time and reached the microsecond level, which indicates that the transition matrix speculated by the molecular simulation of the simulated nanosecond time scale can predict the state transition of the simulated microsecond time scale. From Figure 6a, it can be observed that the implied timescales of the three eigenvalues have converged when the lag time τ=10 ns. Therefore, for the subsequent correctness test (CK test), a lag time of τ=10 ns is used. Figure 6b displays the results of the CK test, with the light blue region representing the confidence interval from multiple tests. The consistency of deviations among multiple tests in Figure 6b is similar to that observed in the dipeptide case, indicating a high level of convergence and prediction time delay in the model. In terms of the probabilistic prediction for the 4-state transition, both curves in the figure closely align, even at 80 ns, demonstrating the excellent prediction accuracy of the model.

**Figure 6.**
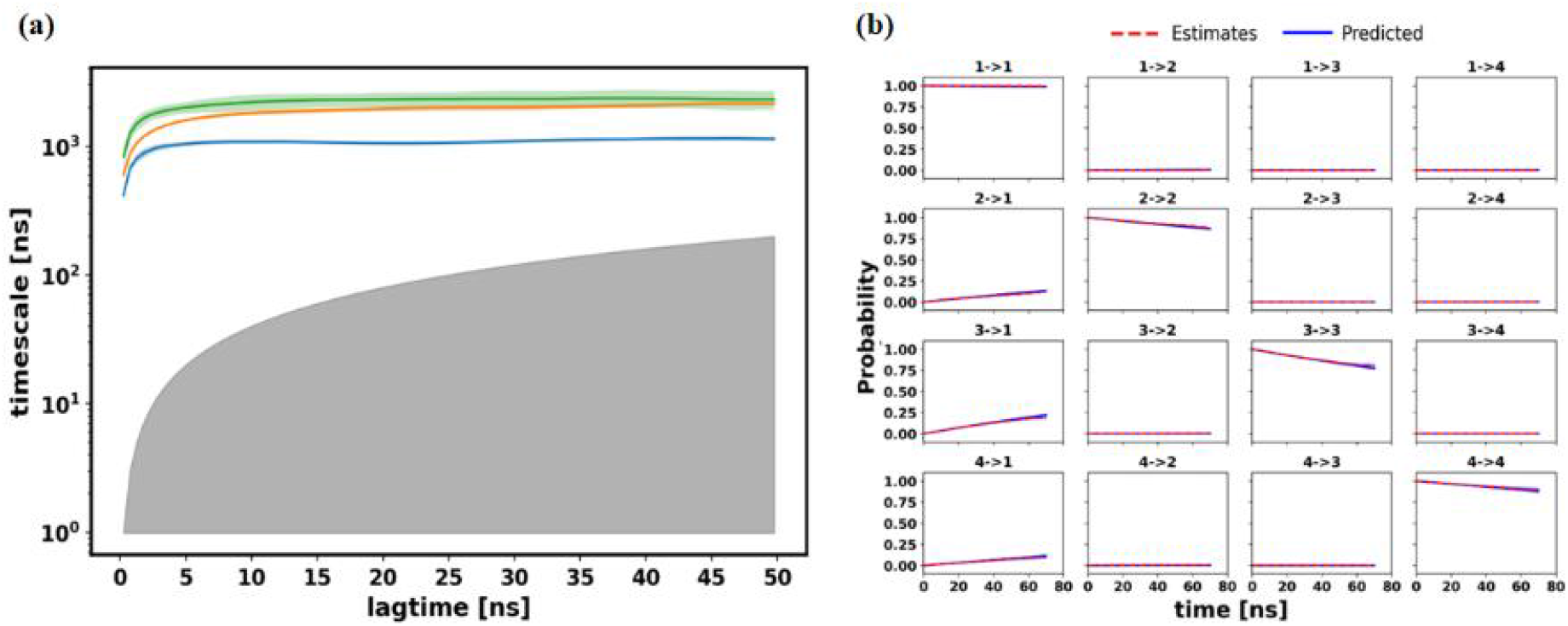
Correctness verification of Aβ_42_. (a) Implied timescale (ITS), (b) CK test results of 6 states.

To compare the impact of adding constraints, the graph embeddings obtained from the model results are projected onto the free energy plane, as depicted in Figure 7. The graph embedding matrix, treated as 2-dimensional data, is utilized to generate a two-dimensional plane using the Pyemma tool. This tool constructs a two-dimensional free energy landscape by plotting a histogram of the scattered data from all trajectories in the training set.

**Figure 7.**
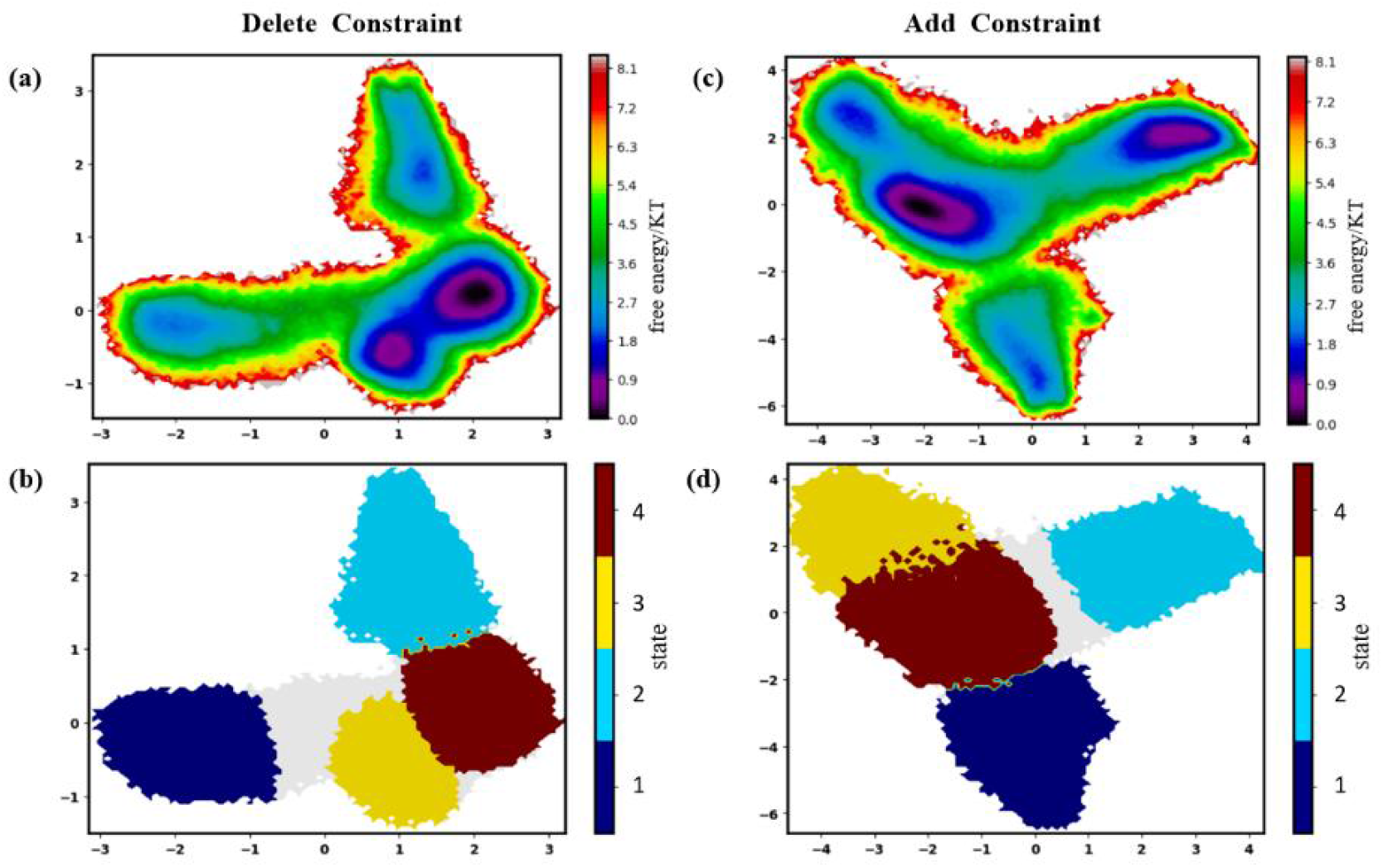
Aβ_42_ free energy surface projection. (a) unconstrained Free energy landscape (FEL) in 2d embedding, and (b) unconstrained classification, (c)constrained Free energy landscape in 2d embedding, and (d) constrained classification.

It is evident from Figure 7c that the model has successfully identified four distinct subgraphs. The protein conformations are accurately partitioned according to their corresponding free energy regions, as observed in subgraph d. It is worth noting that the a and b subgraphs are free energy plane drops without physical constraints. when comparing the model with physical constraints (Figure c) to the unconstrained model (Figure a), it can be observed that the free energy minima of the 3rd and 4th states in Figure a are closely located together, suggesting potential challenges in distinguishing between these two states. Furthermore, the range of the minimum region in the free energy landscape for the first and second states is smaller in Figure a compared to Figure c, indicating improved convergence when constraints are added. This experimentation once again reinforces the significance of incorporating physical constraints in accurately distinguishing transient states.

#### 3.2.2. Interpretability

In order to better explain the results of the model, we visualize the attention parameters of the model into four categories, as shown in Figure 8. The weight distribution observed aligns with the prior knowledge established by previous studies, substantiating the model’s effectiveness. The Aβ_42_ protein is commonly recognized as consisting of three distinct regions. One of the prominent hydrophobic structures is located in the central region, and the strong influence of the 16-22 residues interval shown in Figure 8c can correspond to this hydrophobic region. Furthermore, the presence of an α-helix near the protein end leads to strong interactions between residues, which is verified with the significant weight observed in the 30−36 residues interval in Figure 8c.

**Figure 8.**
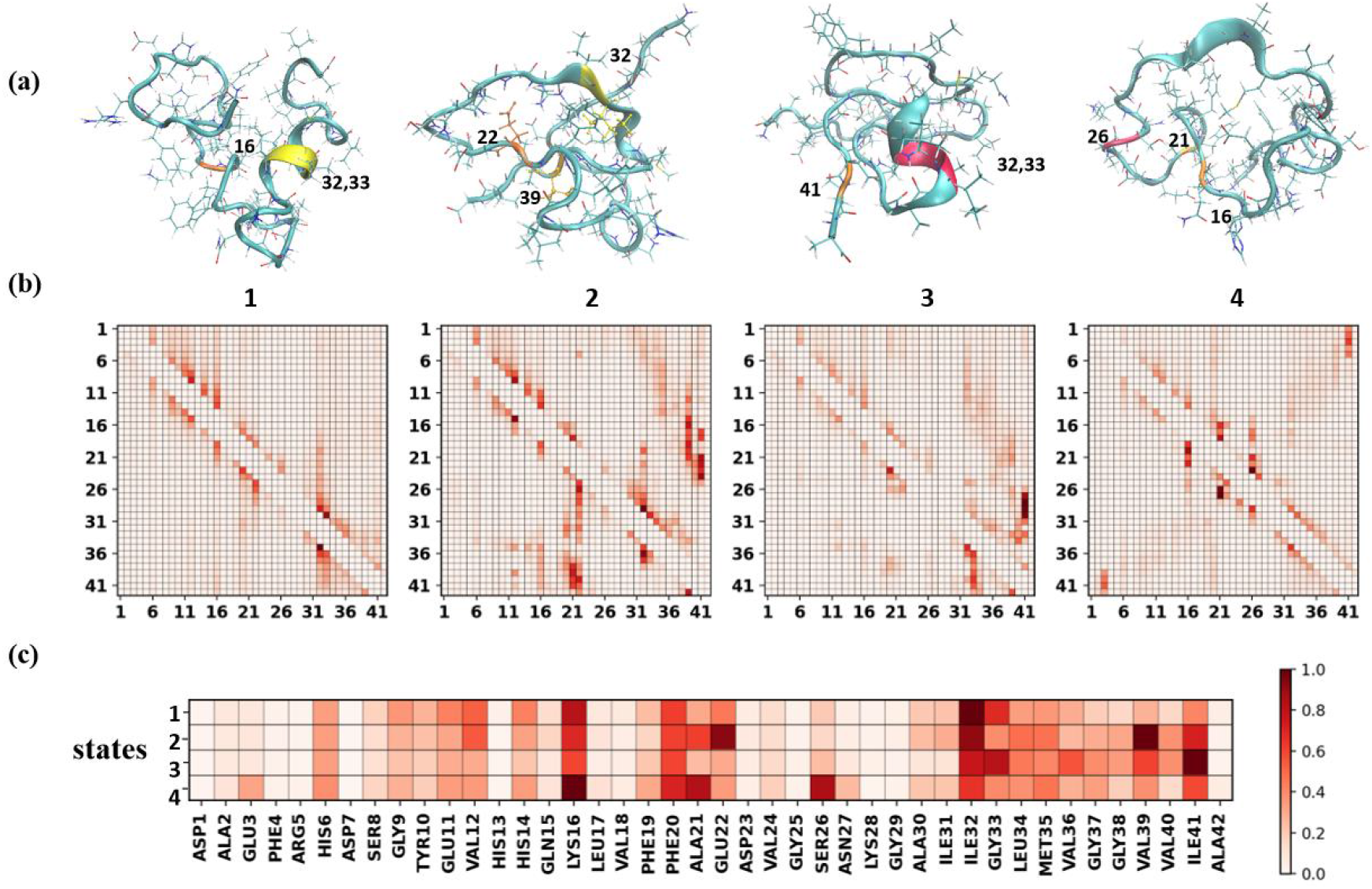
Aβ_42_ visualization results. (a) Typical conformation of 4 states, (b) average attention weights of 42 residue interaction in each state, and (c) average attention weights of 42 residue in different states.

In the first state, GLY33 and ALA30, ILE32 and MET35 play an important role in the interaction in Figure 8b. Figure 8a reveals that these four residues form an α-helix, providing a plausible explanation for their strong interaction. Similarly, in Figure 8c, it can be observed that ILE32 and GLY33 have a higher influence factor in state 1 compared to other states, indicating the crucial role of these four residues specifically in state 1. In state 2, VAL39 with a hydrophobic side chain exhibits a strong influence. It can be seen from Figure a that this residue is relatively close to multiple residues. Moreover, the typical structure of state 2 is surrounded by complex structures, which is also evident in the interaction visualization in Figure 8b. Compared with other states, state 2 exhibits a higher number of significant residues pairs and more complex long-range interactions.

In state 3, the impact of ILE41 stands out as it demonstrates a strong influence on multiple residues, which indicates ILE41 may be a focal point for further research within state 3. Additionally, there is an α-helix in state 3, so ILE32 and GLY33 have important influences. State 4 exhibits a relatively looser structure in Aβ_42_, and the positively charged LYS16 engages in robust ionic interactions. Furthermore, the hydrophobic ALA21 interacts strongly with residues possessing polar side chains such as SER26 and ASN27. These observations highlight the model’s ability to provide biologically meaningful understanding and explanations for corresponding phenomena.

It is worth noting that LYS16 plays an important role in the four states. This charged amino acid usually produces ionic interactions. This strong interaction will have an important impact on the protein structure. Previous studies have indicated the significant involvement of this charged amino acid in the oxidative stress mechanism of Alzheimer’s disease[37], which is consistent with our results. Similarly, ILE32 also plays an important role, owing to its structural position as a constituent residue of the α-helix. In addition, ILE41 and PHE20 with hydrophobic and aromatic side chains are also important in the four states, indicating that these amino acids may be important amino acids of Aβ_42_ protein. This evidence highlights the mutual confirmation between our model results and prior research findings, providing researchers with fresh analytical perspectives and data-supported insights.

## 4. Conclusions

In this paper, a new unsupervised data analysis model of Markov variational method is proposed to realize the accelerated analysis of protein molecular simulation results. The model innovatively combines the learning performance of graph convolutional neural network and physical constraint, and realize the importance modeling of protein molecules through the attention mechanism to provide interpretable molecular mechanisms. By analyzing trajectory data, our approach enhances the understanding of protein targets and enables the extraction of additional information for pharmaceutical researchers, thus fostering advancements in protein research. The graph neural network model in this paper better retains the complex coordinates and structural information in the data in the original simulation, and can model complex multi-body interactions. By incorporating a reversible and balanced physical constraint optimization model during training, the model acquires a robust end-to-end learning capability, enabling the analysis of molecular simulation trajectories containing numerous metastable and transition states. It is beneficial to be applied to natural disordered protein simulations that are difficult for other similar graph-based convolutional neural network models to handle. The VAMP scores of this model in two different scale data sets of alanine dipeptide and Aβ_42_ are higher than those of classic VAMPnets, GCN-VAMP, and MAGNN-VAMP networks. Furthermore, Additionally, we have incorporated an attention mechanism to facilitate interpretability. Through the analysis of attention parameter values, we can assess the intensity of interactions among individual atoms or amino acid residues. This allows us to identify key factors that contribute to protein structure stability and state transitions. Our end-to-end analysis methods provide researchers in related fields with a comprehensive approach that does not rely solely on prior research experience. In the test set, through the visual examination of attention results, we discovered several important amino acids associated with protein state transitions. Remarkably, one of these amino acids was confirmed to be closely linked to Alzheimer’s disease in 2017. By utilizing the visualization of the attention mechanism, this methodology offers protein simulation researchers a novel approach to analysis.

While the method presented in this paper demonstrates promising performance in the experiments, there are still certain challenges that need to be addressed. One such challenge is the impact of the training set on the attention mechanism results. In the case of Aβ_42_ training, the weights associated with residue effects obtained from models trained on different scales of simulated trajectories exhibit subtle variations, with the influence of residues fluctuating within a certain range. This phenomenon may arise from discrepancies in the number of state transitions encompassed by different trajectories. Additionally, the model’s performance is contingent upon the chosen number of states, necessitating manual selection to ensure accurate results, thus prolonging the model training time. In future work, we aim to rectify these issues by refining the input data of the model, enriching its diversity, and further optimizing the network model by incorporating insights from state-of-the-art graph neural models. We aim to offer protein drug researchers an enhanced and dependable tool for molecular simulation analysis, characterized by improved efficiency and reliability.

## Supporting information

support information

## Data availability statement

Code and data will be available at https://github.com/DS00HY/RevGraphVamp upon the publishment of this work but are available from the corresponding author on reasonable request

## Acknowledgements

The authors acknowledge the financial support from National Science Foundation of China (Grant No. 81171488, No. 81671669, No. 81820108018 and No. U22A2041.) and Shenzhen Science and Technology Program (No. KQTD20200820113106007 and No. RCYX20200714114734194). This work is also supported by Shenzhen Key Laboratory of Intelligent Bioinformatics (ZDSYS2 0220422103800001).

## Author contributions statement

H. Y. designed the project, implemented code for all the algorithms and data analysis, and wrote the manuscript. All the authors read and commented on the manuscript and approved the final manuscript.

## Competing interest

The authors declare that they have no competing interests.

## Ethics approval and consent to participate

Not applicable.

## Consent for publication

Not applicable.

## Notes

### Competing Interest Statement

The authors have declared no competing interest.

## Reference

[1] T. Hansson, C. Oostenbrink, W. van Gunsteren, Molecular dynamics simulations, Current opinion in structural biology 12(2) (2002) 190–196.

[2] M. Karplus, J.A. McCammon, Molecular dynamics simulations of biomolecules, Nature structural biology 9(9) (2002) 646–652.

[3] S.A. Hollingsworth, R.O. Dror, Molecular Dynamics Simulation for All, Neuron 99(6) (2018) 1129–1143.

[4] R.O. Dror, R.M. Dirks, J. Grossman, H. Xu, D.E. Shaw, Biomolecular simulation: a computational microscope for molecular biology, Annual review of biophysics 41 (2012) 429–452.

[5] P. Śledź, A. Caflisch, Protein structure-based drug design: from docking to molecular dynamics, Curr Opin Struct Biol 48 (2018) 93–102.

[6] T.I. Adelusi, A.-Q.K. Oyedele, I.D. Boyenle, A.T. Ogunlana, R.O. Adeyemi, C.D. Ukachi, M.O. Idris, O.T. Olaoba, I.O. Adedotun, O.E. Kolawole, Y. Xiaoxing, M. Abdul-Hammed, Molecular modeling in drug discovery, Informatics in Medicine Unlocked 29 (2022) 100880.

[7] X. Gong, Y. Zhang, J. Chen, Advanced Sampling Methods for Multiscale Simulation of Disordered Proteins and Dynamic Interactions, Biomolecules 11(10) (2021) 1416.

[8] F. Chiti, C.M. Dobson, Protein misfolding, functional amyloid, and human disease, Annual review of biochemistry 75 (2006) 333–66.

[9] B.E. Husic, V.S. Pande, Markov state models: From an art to a science, Journal of the American Chemical Society 140(7) (2018) 2386–2396.

[10] T. Hempel, M.J. del Razo, C.T. Lee, B.C. Taylor, R.E. Amaro, F. Noé, Independent Markov decomposition: Toward modeling kinetics of biomolecular complexes, Proceedings of the National Academy of Sciences 118(31) (2021) e2105230118.

[11] B. Liu, Y. Qiu, E.C. Goonetilleke, X. Huang, Kinetic network models to study molecular self-assembly in the wake of machine learning, MRS Bulletin 47(9) (2022) 958–966.

[12] S.-T. Tsai, E.-J. Kuo, P. Tiwary, Learning molecular dynamics with simple language model built upon long short-term memory neural network, Nature Communications 11(1) (2020) 5115.

[13] S. Cao, A. Montoya-Castillo, W. Wang, T.E. Markland, X. Huang, On the advantages of exploiting memory in Markov state models for biomolecular dynamics, The Journal of Chemical Physics 153(1) (2020) 014105.

[14] G. Pérez-Hernández, F. Paul, T. Giorgino, G. De Fabritiis, F. Noé, Identification of slow molecular order parameters for Markov model construction, The Journal of chemical physics 139(1) (2013) 07B604_1.

[15] F. Litzinger, L. Boninsegna, H. Wu, F. Nuske, R. Patel, R. Baraniuk, F. Noé, C. Clementi, Rapid calculation of molecular kinetics using compressed sensing, Journal of Chemical Theory and Computation 14(5) (2018) 2771–2783.

[16] K.A. Konovalov, I.C. Unarta, S. Cao, E.C. Goonetilleke, X. Huang, Markov state models to study the functional dynamics of proteins in the wake of machine learning, JACS Au 1(9) (2021) 1330–1341.

[17] H. Sidky, W. Chen, A.L. Ferguson, High-Resolution Markov State Models for the Dynamics of Trp-Cage Miniprotein Constructed Over Slow Folding Modes Identified by State-Free Reversible VAMPnets, The Journal of Physical Chemistry B 123(38) (2019) 7999–8009.

[18] H. Wu, A. Mardt, L. Pasquali, F. Noe, Deep generative Markov state models, Proceedings of the 32nd International Conference on Neural Information Processing Systems, Curran Associates Inc., Montréal, Canada, 2018, pp. 3979–3988.

[19] H. Wu, F. Noé, Variational Approach for Learning Markov Processes from Time Series Data, Journal of Nonlinear Science 30(1) (2020) 23–66.

[20] A. Mardt, L. Pasquali, H. Wu, F. Noé, VAMPnets for deep learning of molecular kinetics, Nature Communications 9(1) (2018) 5.

[21] W. Chen, H. Sidky, A.L. Ferguson, Nonlinear discovery of slow molecular modes using state-free reversible VAMPnets, The Journal of Chemical Physics 150(21) (2019) 214114.

[22] A. Mardt, L. Pasquali, F. Noé, H. Wu, Deep learning Markov and Koopman models with physical constraints, Mathematical and Scientific Machine Learning, PMLR, 2020, pp. 451–475.

[23] T. Löhr, K. Kohlhoff, G.T. Heller, C. Camilloni, M. Vendruscolo, A kinetic ensemble of the Alzheimer’s Aβ peptide, Nature Computational Science 1(1) (2021) 71–78.

[24] X.-M. Zhang, L. Liang, L. Liu, M.-J. Tang, Graph Neural Networks and Their Current Applications in Bioinformatics, Frontiers in Genetics 12 (2021).

[25] S. Navlakha, C. Kingsford, The power of protein interaction networks for associating genes with diseases, Bioinformatics 26(8) (2010) 1057–1063.

[26] S. Olsson, F. Noé, Dynamic graphical models of molecular kinetics, Proceedings of the National Academy of Sciences 116(30) (2019) 15001–15006.

[27] T. Xie, A. France-Lanord, Y. Wang, Y. Shao-Horn, J.C. Grossman, Graph dynamical networks for unsupervised learning of atomic scale dynamics in materials, Nature Communications 10(1) (2019) 2667.

[28] M. Ghorbani, S. Prasad, J.B. Klauda, B.R. Brooks, GraphVAMPNet, using graph neural networks and variational approach to Markov processes for dynamical modeling of biomolecules, The Journal of Chemical Physics 156(18) (2022) 184103.

[29] K. Schutt, P.-J. Kindermans, H.E. Sauceda Felix, S. Chmiela, A. Tkatchenko, K.-R. Muller, Schnet: A continuous-filter convolutional neural network for modeling quantum interactions, Advances in neural information processing systems 30 (2017).

[30] F. Nuske, H. Wu, J.-H. Prinz, C. Wehmeyer, C. Clementi, F. Noć, Markov state models from short non-equilibrium simulations—Analysis and correction of estimation bias, The Journal of Chemical Physics 146(9) (2017) 094104.

[31] J.C. Stroud, C. Liu, P.K. Teng, D. Eisenberg, Toxic fibrillar oligomers of amyloid-β have cross-β structure, Proc Natl Acad Sci U S A 109(20) (2012) 7717–22.

[32] G.F. Chen, T.H. Xu, Y. Yan, Y.R. Zhou, Y. Jiang, K. Melcher, H.E. Xu, Amyloid beta: structure, biology and structure-based therapeutic development, Acta Pharmacol Sin 38(9) (2017) 1205–1235.

[33] W.C. Swope, J.W. Pitera, F. Suits, Describing Protein Folding Kinetics by Molecular Dynamics Simulations. 1. Theory, The Journal of Physical Chemistry B 108(21) (2004) 6571–6581.

[34] M. Hoffmann, M. Scherer, T. Hempel, A. Mardt, B. de Silva, B.E. Husic, S. Klus, H. Wu, N. Kutz, S.L. Brunton, F. Noé, Deeptime: a Python library for machine learning dynamical models from time series data, Machine Learning: Science and Technology 3(1) (2022) 015009.

[35] M.K. Scherer, B. Trendelkamp-Schroer, F. Paul, G. Pérez-Hernández, M. Hoffmann, N. Plattner, C. Wehmeyer, J.-H. Prinz, F. Noé, PyEMMA 2: A Software Package for Estimation, Validation, and Analysis of Markov Models, Journal of Chemical Theory and Computation 11(11) (2015) 5525–5542.

[36] A. Vaswani, N. Shazeer, N. Parmar, J. Uszkoreit, L. Jones, A.N. Gomez, L. Kaiser, I. Polosukhin, Attention is all you need, Advances in neural information processing systems 30 (2017).

[37] S.M. Fica-Contreras, S.O. Shuster, N.D. Durfee, G.J.K. Bowe, N.J. Henning, S.A. Hill, G.D. Vrla, D.R. Stillman, K.M. Suralik, R.K. Sandwick, S. Choi, Glycation of Lys-16 and Arg-5 in amyloid-β and the presence of Cu(2+) play a major role in the oxidative stress mechanism of Alzheimer’s disease, J Biol Inorg Chem 22(8) (2017) 1211–1222.

