## Supplementary material for "RevGraphVAMP: A protein molecular simulation analysis model combining graph convolutional neural networks and physical constraints": support information

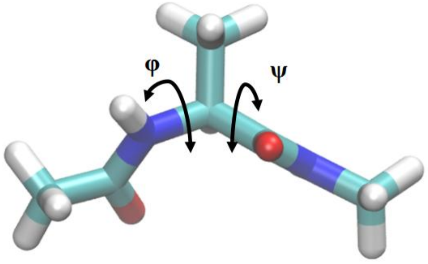


Figure S1. Alanine dipeptide

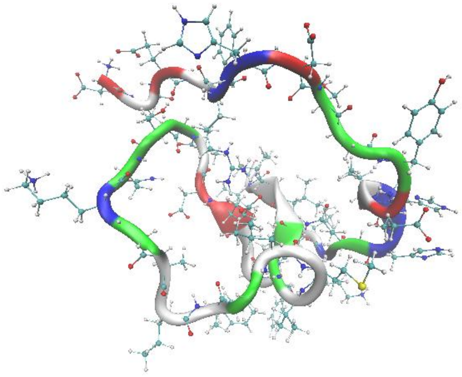

Figure S2. $A\beta_{42}$ structure

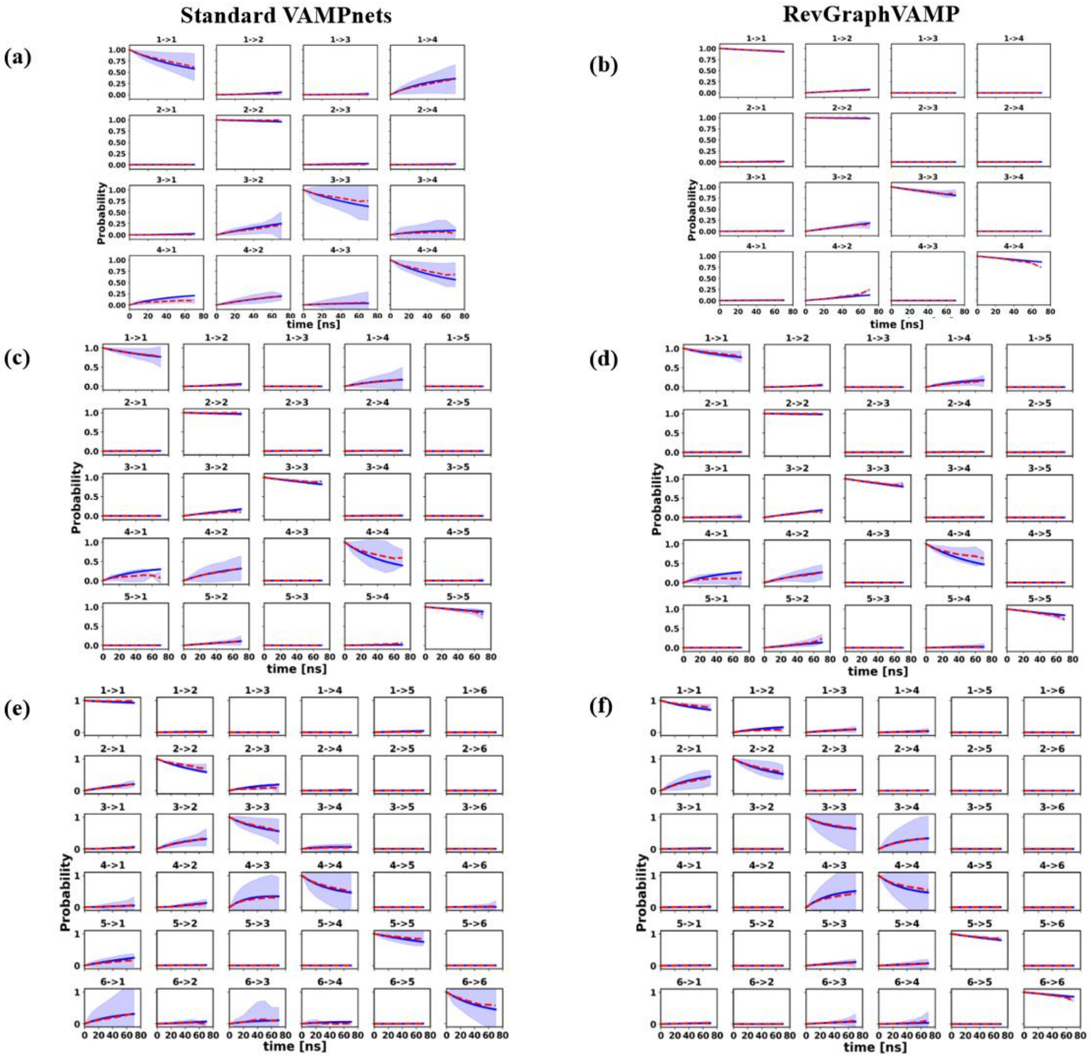


**Figure S3.** Aβ42 protein CK test results comparison of Standard VAMPnets and RevGraphVAMP at $\tau = 10$ ns. The horizontal coordinate is the predicted step length, the vertical coordinate is the transfer probability, the blue solid line is the estimated value K(τ) of the transfer probability under this time, the red part is the predicted value K(nτ) calculated by using the K matrix according to Eq.(28), the light blue part is the confidence interval of multiple tests. (a), (c) and (e) on the left are the prediction results of standard VAMPnets. (b), (d) and (e) on the right are the prediction results of our model.

**Table S1**. The input and type of the model

| Model | Input | Net of $\chi(x)$ | Reference |
|---|---|---|---|
| **VAMPnets** | Alanine dipeptide molecular coordinates (30 dimensions) | Fully-connected network | [1] |
| | Distance between $A\beta_{42}$ residues (710 dimensions) | Fully-connected network | [2] |

| | | | |
|---|---|---|---|
| **GCN-VAMP** | Graph data feature | Graph convolutional network | [3] |
| **MAGNN-VAMP** | Graph data feature | Multi-head attention Graph network | - |
| **RevGraphVAMP** | Graph data feature | GASchNet | - |

Table S2. The proportion of each state of alanine dipeptide

| **Model** | **1** | **2** | **3** | **4** | **5** | **6** |
|---|---|---|---|---|---|---|
| **VAMPnets** | 39.47% | 23.52% | 23.16% | 10.60% | 3.07% | 0.15% |
| **RevGraphVAMP** | 39.69% | 24.67% | 22.55% | 10.10% | 2.84% | 0.15% |

### Reference


[1] A. Mardt, L. Pasquali, H. Wu, and F. Noé, “VAMPnets for deep learning of molecular kinetics,” *Nature Communications,* vol. 9, no. 1, pp. 5, 2018/01/02, 2018.

[2] T. Löhr, K. Kohlhoff, G. T. Heller, C. Camilloni, and M. Vendruscolo, “A kinetic ensemble of the Alzheimer’s Aβ peptide,” *Nature Computational Science,* vol. 1, no. 1, pp. 71-78, 2021.

[3] T. Xie, A. France-Lanord, Y. Wang, Y. Shao-Horn, and J. C. Grossman, “Graph dynamical networks for unsupervised learning of atomic scale dynamics in materials,” *Nature Communications,* vol. 10, no. 1, pp. 2667, 2019/06/17, 2019.
